# Screening of FDA-approved drugs using a MERS-CoV clinical isolate from South Korea identifies potential therapeutic options for COVID-19

**DOI:** 10.1101/2020.02.25.965582

**Authors:** Meehyun Ko, So Young Chang, Soo Young Byun, Aleksandr Ianevski, Inhee Choi, Anne-Laure Pham Hung d’Alexandry d’Orengiani, Denis E. Kainov, David Shum, Ji-Young Min, Marc P. Windisch

## Abstract

Therapeutic options for coronavirus remain limited. To address this unmet medical need, we screened 5,406 compounds, including United States Food and Drug Administration (FDA)- approved drugs and bioactives, for activity against a South Korean Middle East respiratory syndrome coronavirus (MERS-CoV) clinical isolate. Among 221 identified hits, 54 had therapeutic indexes (TI) greater than 6. Time-of-addition studies with selected drugs demonstrated eight and four FDA-approved drugs acted on the early and late stages of the viral life cycle, respectively. Confirmed hits included several cardiotonic agents (TI>100), atovaquone, an anti-malarial (TI>34), and ciclosonide, an inhalable corticosteroid (TI>6). Furthermore, utilizing the severe acute respiratory syndrome CoV-2 (SARS-CoV-2), combinations of remedesivir with selected dugs were evaluated, which identified ciclosonide and nelfinavir to be additive and synergistic drugs *in vitro*, respectively. Together, we screened FDA-approved drugs using patient-derived MERS-CoV, triaged hits to discriminate between early and late viral life cycle inhibitors, confirmed selected drugs using SARS-CoV-2, and demonstrated the added value of selected medications in combination with remedesivir. Our results identify potential therapeutic options for MERS-CoV infections, and provide a basis to treat coronavirus disease 2019 (COVID-19) and other coronavirus-related illnesses.

## 1. Introduction

Coronaviruses (CoV) are enveloped, positive-sense, single-stranded RNA viruses in the *Coronaviridae* family of the *Nidovirales* order. CoV usually cause mild to severe respiratory tract infections [1]. The two types of human coronaviruses that had been described prior to 2003, coronavirus 229E and OC43, caused mild, cold-like symptoms [2,3]. However, an outbreak of Severe Acute Respiratory Syndrome (SARS) in 2003, which occurred mainly in Southeast Asia, was attributed to a coronavirus. That outbreak resulted in 8,096 confirmed cases and 774 deaths (fatality rate of 9.6%) [4].

In 2012, the novel coronavirus Human Coronavirus-Erasmus Medical Center (HCoV-EMC) was isolated from a patient in Saudi Arabia who developed pneumonia and renal failure [5]. From the first outbreak in 2012 until January 2019, the HCoV-EMC epidemic resulted in 2,449 laboratory-confirmed cases and at least 845 deaths (fatality rate 34%), mainly in the Arabian Peninsula. Thus, HCoV-EMC was renamed Middle East respiratory syndrome coronavirus (MERS-CoV) [6]. Another major outbreak of MERS-CoV infection, the largest outside the Arabian Peninsula, occurred in South Korea in 2015 [7,8]. Notably, aside from the index case of MERS-CoV, the majority of viral transmissions in South Korea were nosocomial, with 186 confirmed cases across 16 clinics [7,9]. Furthermore, the World Health Organization (WHO) has reported continual waves of MERS outbreaks in the Middle East, although they have been smaller than the major 2014 outbreak [6].

Due to the severity of MERS infection and the urgent need for effective treatment, several approaches for therapeutic development have been attempted [10]. In clinical studies, a combination of ribavirin and interferon-alpha (IFN-α) therapy improved patient survival rates when administered early after the onset of infection, but had no significant effect in the late stage of infection [11-13]. These results suggest that broad-spectrum antivirals can be effective in MERS patients at some stages of infection, but for complete antiviral activity, a treatment specific for MERS-CoV may be required.

With the recent emergence of the severe acute respiratory syndrome CoV-2 in Wuhan, China in late 2019, coronavirus disease 2019 (COVID-19) is rapidly spreading worldwide. The ongoing COVID-19 pandemic has already caused many human casualties and significant socio-economic losses globally. With more than 60 million COVID-19 cases confirmed and over 1.4 million related fatalities reported (November 26^th^, 2020), there is worldwide effort to control the spread of this devastating virus. Unfortunately, there are no CoV-specific drugs approved by the United States Food and Drug Administration (FDA) for clinical use, nor repurposed drugs to efficiently treat COVID-19 patients, yet.

## 2. Materials and Methods

### 2.1 Cell line and virus

Vero cells (ATCC CCL-81; Manassas, VA, USA) were maintained at 37°C with 5% CO_2_ in Dulbecco’s Modified Eagle’s Medium (Welgene, Gyeongsan, Republic of Korea) supplemented with 10% heat-inactivated fetal bovine serum and 1X antibiotic-antimycotic solution (Gibco/Thermo Fisher Scientific, Waltham, MA, USA). Calu-3 cells (ATCC HTB-55; Manassas, VA, USA) were maintained at 37°C with 5% CO_2_ in Eagle’s Minimum Essential Medium (ATCC, Manassas, VA, USA) with 10% heat-inactivated fetal bovine serum and 1X antibiotic-antimycotic solution (Gibco/Thermo Fisher Scientific, Waltham, MA, USA). The Korean strain of MERS-CoV (MERS-CoV/KOR/KNIH/002_05_2015; MERS/KOR/2015, Genbank accession no. KT029139.1) [14] was kindly provided by Sung Soon Kim, Division of Respiratory Viruses, Center for Infectious Diseases, Korea National Institute of Health (KNIH), Korea Centers for Disease Control and Prevention (KCDC), and propagated in Vero cells, as previously described [15]. Viral titers were determined by plaque assays in Vero cells as described [16]. All experiments using MERS-CoV were performed at Institut Pasteur Korea in compliance with the guidelines of the KNIH using enhanced Biosafety Level 3 (BSL-3) containment procedures in laboratories approved for use by the KCDC. The SARS-CoV-2 hCoV-19/Norway/Trondheim-S15/2020 strain has been described in a previous study [17]. Briefly, the virus was amplified in Vero-E6 cells incubated in DMEM containing Pen/Strep and 0.2% bovine serum albumin. All experiments using SARS-CoV-2 were performed in the BSL-3 at NTNU.

### 2.2 Compound libraries

A compound library of 5,406 compounds composed of FDA-approved drugs, which covers approximately 60% of all FDA-approved compounds, bioactives, kinase inhibitors, and natural products, was compiled (LOPAC, Prestwick, Microsource, Selleck, Tocris) and used for this screen. Compounds were dissolved in DMSO at 10 mM and stored at −80°C until use.

### 2.3 Image-based screening assay and assay validation

Vero cells were seeded at 1.2 × 10^4^ cells per well in Opti-PRO™ SFM supplemented with 4 mM L-glutamine and 1X Antibiotic-Antimycotic solution (Gibco/Thermo Fisher Scientific) in black, 384-well, μClear plates (Greiner Bio-One, Kremsmünster, Austria) at 24 h prior to the experiment. Subsequently, compounds were added to each well using an automated liquid handling system (Apricot Designs, Covina, CA, USA) before virus infection. Final concentrations of each compound were 10 μM, and the DMSO concentration was kept at 0.5% or lower. For viral infection, plates were transferred into the BL-3 containment facility to add MERS-CoV at a multiplicity of infection (MOI) of 0.0625, and cells were fixed at 24 hours post-infection (hpi) with 4% PFA followed by immunofluorescence analyses. MERS-CoV infection was detected using rabbit anti-MERS-CoV S antibodies, and cell viability was evaluated by Hoechst 33342 staining. Images were acquired by a Perkin Elmer Operetta (20×; Waltham, MA, USA) and analyzed by in-house developed Image Mining 3.0 (IM 3.0) plug-in software. To validate the assay, dose-response curves (DRCs) with four compounds with known antiviral activities against MERS-CoV were assessed: CQ, CPZ, CsA, and LPV [18,19]. Compounds with >70% MERS-CoV inhibition and >70% viability were subjected to DRC analyses, as described below.

### 2.4 Dose-response curve drug analysis

The primary hits (256 hits) were used to generate 10-point DRCs, with compound concentrations from 0.05–25 μM. The acquired images were analyzed using in-house software to quantify cell numbers and infection ratios. The antiviral activity was normalized to positive (mock) and negative (0.5% DMSO) controls in each assay plate. DRCs were fitted by sigmoidal dose-response models, and the equation was described as Y = Bottom + (Top - Bottom)/(1 + (IC_50_/X)^Hillslope^) using XLfit 4 Software or Prism7. The IC_50_ was calculated from the normalized activity data set fitted curve. All IC_50_ and CC_50_ values were measured in duplicate and the quality of each assay was controlled by Z′-factor and the coefficient of variation in percent (%CV).

### 2.5 Pharmacological action clustering

The information regarding pharmacological actions of each compound was compiled by using ChemIDPlus and MESH databases [20,21] and information provided by the vendors. Once relevant information was collected, pharmacological actions were manually reassessed to finally categorize all compounds into 43 different pharmacological actions. The information on approval status for drugs was retrieved from DrugBank, version 5.0.7 [22].

### 2.6 Drug combination studies

Vero-E6 cells were treated with different concentrations of two drugs and infected with SARS-CoV-2 at an MOI 0.1 or mock. At 72 hpi, cell viability was measured using CellTiter-Glo (Promega, Madison, WI, USA). The observed responses were compared to expected combination responses. The expected responses were calculated based on the zero interaction potency (ZIP) reference model using SynergyFinder version 2 [17]. Briefly, the degree of combination synergy, or antagonism, is quantified by comparing the observed drug combination response against the expected response, calculated using a reference model that assumes no interaction between drugs. Synergy scores, which represent an averaged percentage excess effect due to interactions between drugs and are the most common metric to summarize the drug interaction, were quantified. Furthermore, the cytotoxicity of each drug combination, determined by the quantification of ATP level (CellTiter-Glo), was subtracted. Combinations with scores >10 are considered synergistic, scores between −10 and 10 additive, and below −10 antagonistic

## 3. Results

To address the urgent unmet need to develop effective treatments for CoV patients, we implemented a high-content screening (HCS) strategy with the goal of repurposing newly identified MERS-CoV inhibitors for a wider range of CoV, including COVID-19. Utilizing a Korean MERS-CoV patient isolate, we screened 5,406 compounds, including FDA-approved drugs, bioactive agents, kinase inhibitors, and natural products. Our library included 60% of all FDA-approved drugs (1,247 out of 2,069 total) (Fig. 1A). Compounds were tested for activity against MERS-CoV by analyzing the levels of expression of viral spike (S) protein in infected Vero cells using immunofluorescence analysis (IFA). The screens included the reference inhibitor chloroquine (IC_90_ = 93 μM) at 100 μM to define maximum inhibition (De Wilde et al., 2014). The calculated Z’-factor above 0.78 indicated good discrimination between the control dimethyl sulfoxide (DMSO) and chloroquine treatment of infected cells (Fig. 1B). Two independent HCS analyses (screen 1 and screen 2) were conducted, demonstrating a high degree of correlation (R^2^ = 0.91) between the two replicates (Fig. 1C). These screens identified 256 compounds that demonstrated >70% MERS-CoV inhibition and >70% cell viability (Fig. 1D). These primary hits were then confirmed using 10-point dose-response curve (DRC) analysis to determine IC_50_ and 50% cytotoxicity concentrations (CC_50_) for each compound (Fig. 1D). A representative DRC analysis is shown in Supplementary Fig. 1. The therapeutic index (TI) was calculated as the ratio of CC_50_/IC_50_. Among the 256 initial hits, 35 compounds were denoted as inactive (TI values <1), and were eliminated from the list of confirmed hits. Of the resulting 221 confirmed hits, 54 compounds with an *in vitro* TI >6 were selected for further testing (Fig. 1D).

**Fig. 1.**
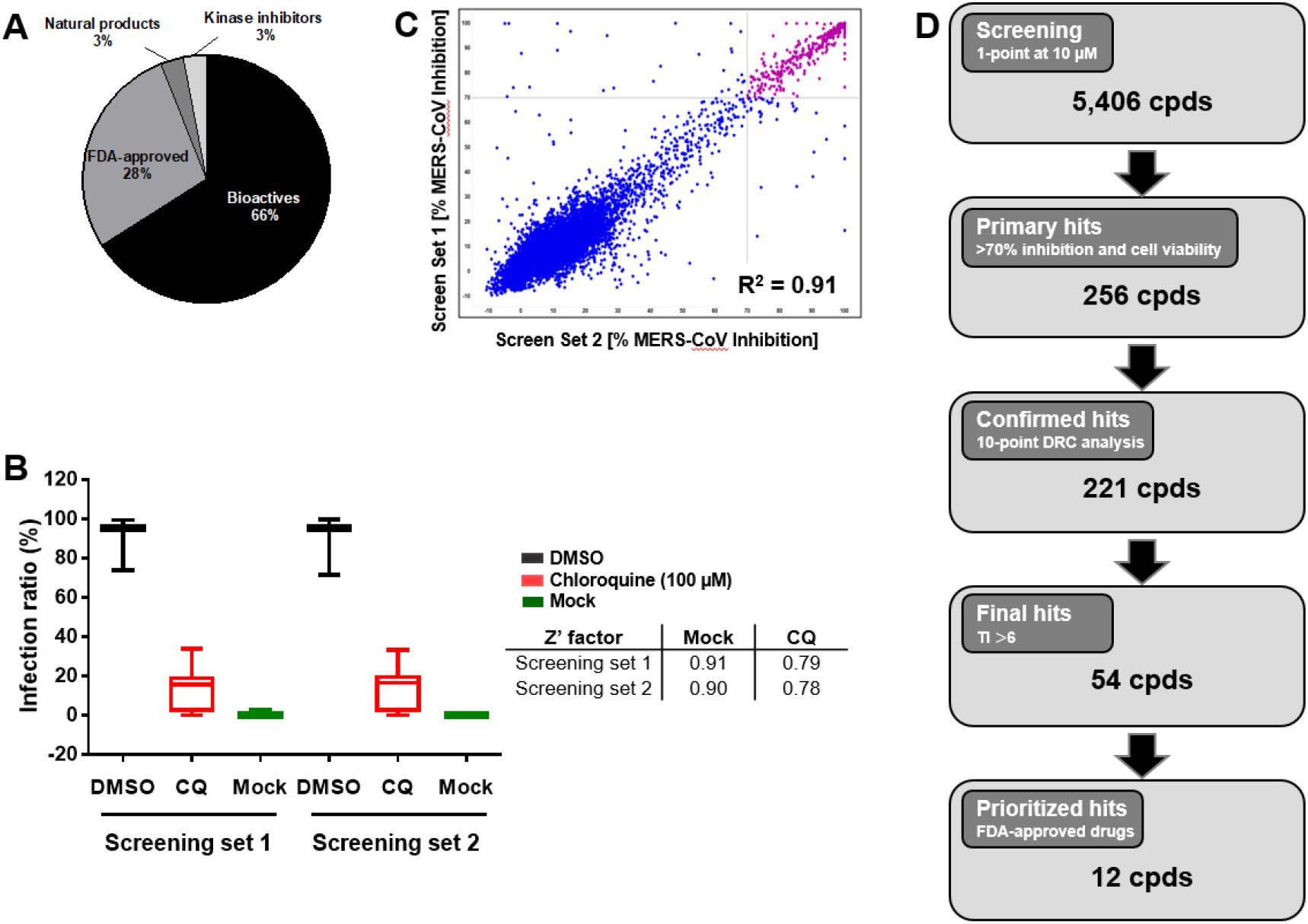
Overview of library composition and triage of hits. (A) Our small-molecule compound library primarily comprised bioactives and FDA-approved drugs, with a small proportion of natural products and kinase inhibitors. (B) High-content screening (HCS) of 5,406 compounds (cpds) in two batches in duplicate, and calculation of Z’-factors between high (MERS-CoV infection, black) and low (mock, green) values. (C) Correlation between duplicate screens. The scatter plot shows MERS-CoV inhibition ratios overlaid with cell viability ratios. Compounds with MERS-CoV inhibition >70% and cell viability >70% were regarded as primary hits. (D) Flowchart of HCS hit selection and confirmation of final hit selection.

To investigate whether the FDA-approved drugs act on the early or late stages of the viral life cycle (pre- or post-entry), we conducted time-of-addition studies. Vero cells were treated with each drug at a concentration above its IC_90_, and analyzed as described in the Supplementary Information. Chloroquine served as an early-stage inhibitor control, and inhibited MERS-CoV infection by up to 30% until 3 hpi. However, chloroquine had no significant effect when administered at 4 hpi (Fig. 2). A similar outcome was observed for treatment with ouabain, digitoxin, digoxin, niclosamide, regorafenib, nelfinavir mesylate, ciclesonide, and benidipine hydrochloride, all of which inhibited MERS-CoV infection only when administered earlier than 4 hpi (Fig. 2, Supplementary Fig. 2). In contrast, atovaquone, lercanidipine hydrochloride, permethrin, and octocrylene had only minor inhibitory effects throughout the time-course assay (Supplementary Fig. 2).

**Fig. 2.**
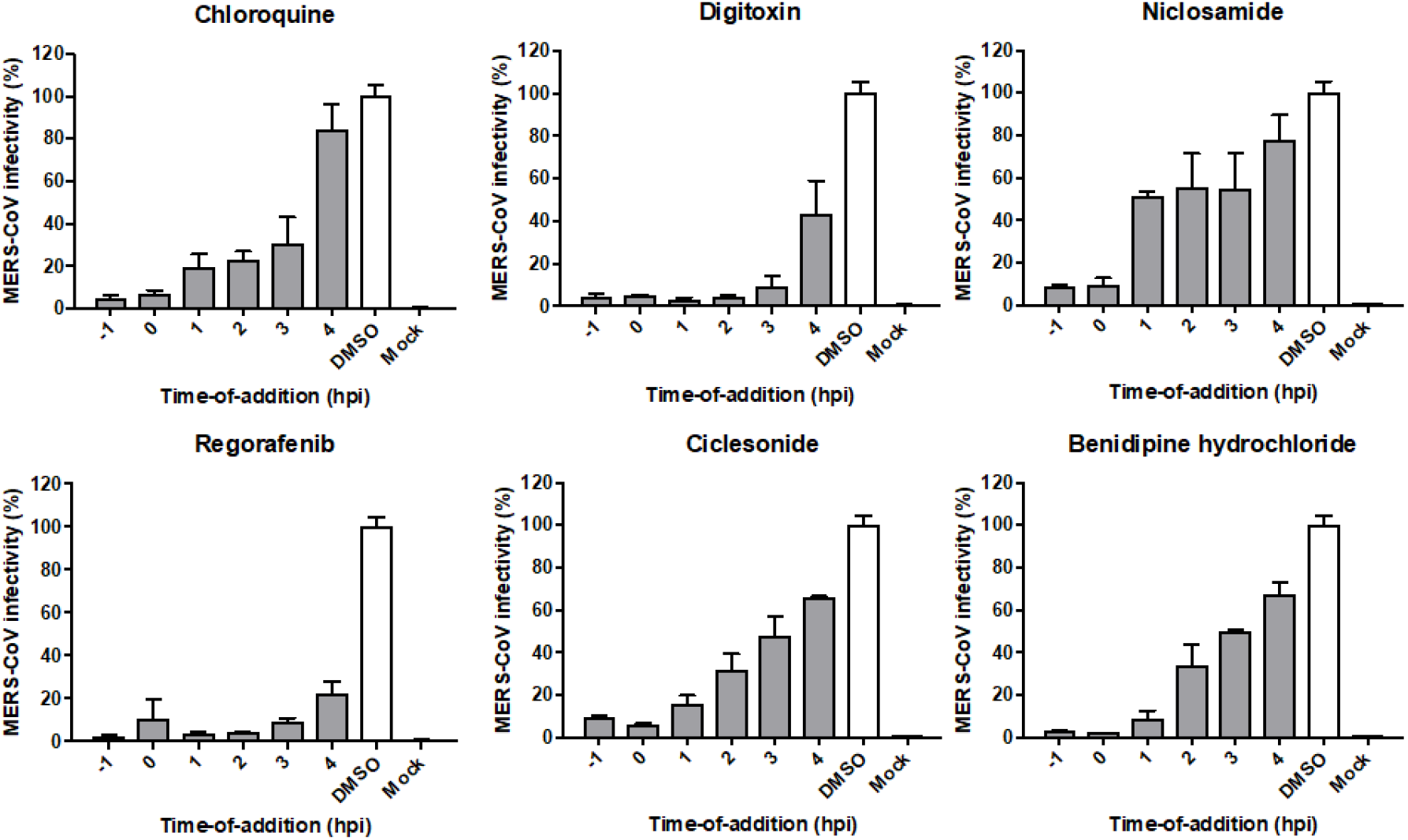
Time-of-addition study with selected FDA-approved drugs. Five FDA-approved drugs were analyzed by time-course experiments to determine the stage of the MERS-CoV life cycle inhibited. Vero cells were infected with MERS-CoV at a multiplicity of infection of 5, and FDA-approved drugs were administered at six time points pre- or post-infection as indicated. Drugs were used at concentrations above their 90% inhibitory concentration (IC_90_) values. Chloroquine served as a known early-stage inhibitor.

Remdesivir, a drug which was originally developed for the treatment of Ebola virus, was recently approved for the treatment of SARS-CoV-2 infection. In previous experiments, we uncovered synergism between remedesivir and several other drugs against SARS-CoV-2 in Calu-3 cells [17]. Therefore, we tested remdesivir in combination with nelfinavir or ciclesonide. As references, we used camostat and cepharanthine in SARS-CoV-2- and mock-infected Vero-E6 cells, and evaluated virus-mediated cytotoxicity by determining ATP level as described before. Each drug combination was tested by a 6×6 dose-response matrix, where five doses of single drugs are combined in a pairwise manner. We subtracted drug combination responses measured on virus-infected cells from those measured on mock-infected cells. As a result, we obtained dose-response matrices demonstrating selective virus inhibition achieved by each combination. For each drug combination, we calculated ZIP synergy scores and plotted synergy distribution maps, showing synergy at each pairwise dose. Thereby, we observed additive effects between remdesivir and cepharanthine, as well as remdesivir and ciclesonide, whereas remdesivir-camostat and remdesivir-nelfinavir combinations showed synergistic effects (Fig. 3A-D). Drug combination results are summarized in Table 1. Thus, these drug combinations should be further investigated *in vitro* and *in vivo*.

**Table 1.** Drug combinations in SARS-CoV-2 infected Vero cells.

| Drug 1 | Drug 2 | ZIP score<sup>1</sup> | Description |
| --- | --- | --- | --- |
| Camostat | Remdesivir | 21.3 | Synergy |
| Nelfinavir | Remdesivir | 13.9 | Synergy |
| Cepharathine | Remdesivir | 4.2 | Additive |
| Ciclesonide | Remdesivir | -0.6 | Additive |

<sup>1</sup>https://synergyfinder.fimm.fi

**Fig. 3.**
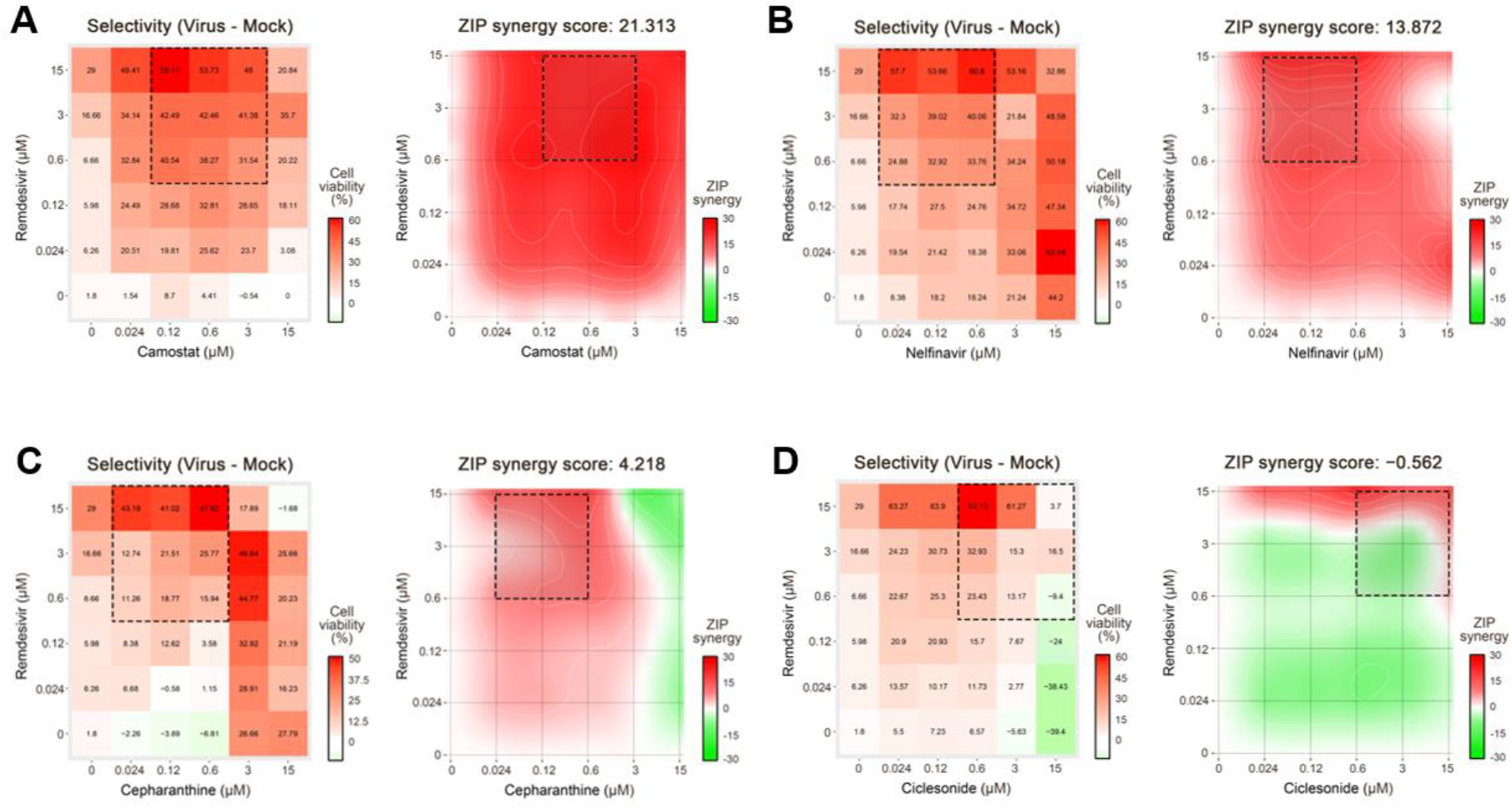
Drug combination studies. Evaluation of drug combinations in SARS-CoV-2 infected Vero-E6 cells. As a read-out, virus-mediated cell death in the presence and absence of drugs was assessed. (A) Remdesivir-camostat, (B) remdesivir-nelfinavir, (C) remdesivir-cepharanthine, and (D) remdesivir-ciclesonide combinations were monitored. Dose-response matrices and synergy distribution maps are shown on the right and left panels with corresponding cell viability and ZIP synergy, respectively. X- and y-axes indicate drug concentrations (μM). ZIP synergy scores were calculated as described in the Material and Methods section.

## Discussion

Our approach aimed to identify FDA-approved drugs and bioactives that could be promptly repurposed or developed, respectively, to treat MERS-CoV and potentially COVID-19-infected patients. In previously reported studies, small molecule libraries that were screened against MERS-CoV included approximately 300 drugs with FDA approval or that were in clinical development [19,23]. Our screen included 1,247 FDA-approved drugs, and as a result, we identified drugs not found in previous studies, indicating that further opportunities exist for identifying novel anti-CoV drugs by screening larger libraries of FDA-approved drugs and bioactives. Moreover, despite having used a different viral isolate than in earlier reports, we corroborated four previously identified hits, including emetine dihydrochloride, ouabain, cycloheximide, and nelfinavir mesylate. This strongly suggests that the drugs reproducibly identified in our HCS assays and in the previously published screens could be repurposed as potential therapeutic options for patients suffering from CoV infections [23].

Fig. 4 shows classification of library compounds into 43 categories of pharmacological action, according to publicly available drug databases. Notably, the cardiovascular agents category contained 14 of the 54 final hit compounds (26%). These belong to a class of cardiac glycosides, naturally derived agents that are used for treating cardiac abnormalities and modulating sodium-potassium pump action [24]. Glycosides have also been reported to exhibit antiviral activity against the herpes simplex virus and human cytomegalovirus [25,26]. Consistent with these previous studies, our data indicate that the cardiac glycosides ouabain, digitoxin, and digoxin also efficiently inhibit MERS-CoV infection. Ouabain has been found to block cellular entry by CoV, such as MERS-CoV, through Src kinase signaling [27]. Based on these data, we speculate that cardiac glycosides may exert anti-MERS-CoV activity through blockade of viral entry. However, more experimental work will be required to elucidate the exact mechanism by which this occurs.

**Fig. 4.**
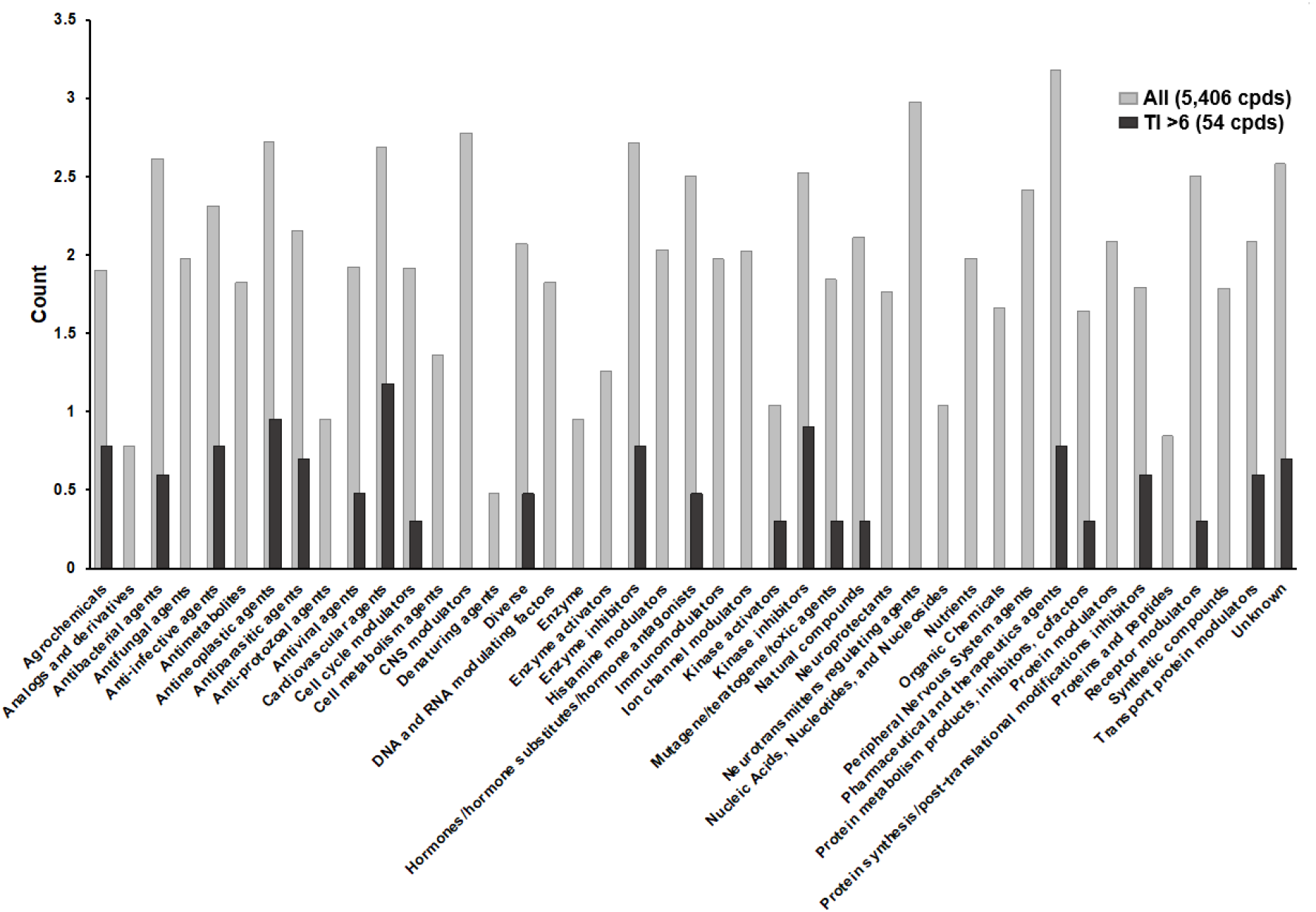
Pharmacological action profiling of all library compounds and confirmed hits. The 54 final hits were sorted into 43 pharmacological action categories. Gray and black bars indicate the distribution of all screened compounds and confirmed hits with a therapeutic index (TI) >6. The vertical axis displays counts of each compound on a log scale.

Drug development may be hastened by repurposing FDA-approved drugs and inhibitors with known biological functions, pharmacological activities, and safety profiles. Therefore, we prioritized 12 FDA-approved drugs and six bioactives not yet reported to have anti-CoV activities; their information is summarized in Tables 2 and 3, respectively. Important to note, a follow-up study confirmed seven of the 12 FDA-approved drugs listed in Table 1 as active against SARS-CoV-2 [28]. An additional 26 inhibitors that our HCS identified include bioactives and drugs that have been studied in clinical trials. A ranking of these inhibitors according to therapeutic index (SI) values, ranging from >6 to >156, is shown in Supplementary Table 1.

**Table 2.** Hit profiling and anti-MERS-CoV efficacies of FDA-approved drugs in Vero cells.^1^.

| Drug Name | Trade Name | Putative Drug Target | Pharmaceutical Action | IC<sub>50</sub><sup>2</sup> (μM) | SD<sup>3</sup> (±) | CC<sub>50</sub><sup>4</sup> (μM) | TI<sup>5</sup> |
| --- | --- | --- | --- | --- | --- | --- | --- |
| Ouabain<sup>#,†</sup> | Strodival | Na, K-exchanging ATPase pump | Cardiotonic agent | 0.08 | 0.0066 | >25<sup>§</sup> | >312.5 |
| Digitoxin<sup>#,†</sup> | Digitaline | Ca, Na-exchanging ATPase pump | Cardiotonic agent | 0.16 | 0.0003 | >25<sup>§</sup> | >156.3 |
| Digoxin<sup>#,†</sup> | Lanoxin | Ca, Na-exchanging ATPase pump | Cardiotonic agent | 0.17 | 0.0084 | >25<sup>§</sup> | >147.1 |
| Niclosamide<sup>#,†</sup> | Niclocide, others | ATP synthase | Agrochemical | 0.55 | 0.0363 | >25<sup>§</sup> | >45.5 |
| Atovaquone* | Mepron | unknown (lipophilic) | Anti-infective agent | 0.72 | 0.0585 | >25 | >34.7 |
| Regorafenib<sup>#,†</sup> (Bay 73-4506) | Stivarga | Multiple kinases | Anti-neoplastic agent | 2.31 | 0.0834 | >25 | >10.8 |
| Lercanidipine hydrochloride* | Zanidip | Calcium channel blocker | Cardiovascular agent | 2.36 | 0.1654 | >25 | >10.6 |
| Permethrin* | Elimite, others | Na channel | Agrochemical | 3.60 | 0.7573 | >25 | >6.9 |
| Octocrylene* | None | Skin | Additive in sunscreen | 3.62 | 0.6435 | >25 | >6.9 |
| Nelfinavir mesylate<sup>#,†</sup> | Viracept | HIV-1 protease | Antiviral agent | 3.62 | 0.0177 | >25 | >6.9 |
| Ciclesonide<sup>#,†</sup> | Alvesco, others | Glucocorticoid ligand | Pharmaceutical and therapeutic agents | 4.07 | 0.4907 | >25<sup>§</sup> | >6.1 |
| Benidipine hydrochloride<sup>#</sup> | Coniel | Calcium channel blocker | Cardiovascular agent | 4.07 | 0.7234 | >25 | >6.1 |

<sup>1</sup>DrugBank database (version 5.0) was used for characterizing FDA-approved drugs
<sup>2</sup>50% inhibitory concentration (IC<sub>50</sub>)
<sup>3</sup>Standard deviation (SD) of replicated IC<sub>50</sub> values <sup>4</sup>50% cytotoxicity concentration (CC<sub>50</sub>) <sup>5</sup>Therapeutic Index (TI): ratio of CC<sub>50</sub>/IC<sub>50</sub> <sup>#</sup>Drug acting on the early stage of the viral life cycle, according to time-of-addition study <sup>\*</sup>Drug acting on the late stage of the viral life cycle, according to time-of-addition study <sup>†</sup>Activity in SARS-CoV-2 system [28] <sup>§</sup>CC<sub>50</sub> >50 µM in Vero cells [28]

**Table 3.** Hit profiling and anti-MERS-CoV efficacies of selected bioactives in Vero cells.^1^.

| Inhibitor Name | Pharmaceutical Action | IC<sub>50</sub><sup>2</sup><br>(µM) | SD<sup>3</sup><br>(±) | CC<sub>50</sub><sup>4</sup><br>(µM) | TI<sup>5</sup> |
| --- | --- | --- | --- | --- | --- |
| Emetine dihydrochloride | Anti-neoplastic agent | 0.08 | 0.0054 | >25 | >312.5 |
| Oxyclozanide | Antiparasitic agent | 0.07 | 0.0060 | 20.92 | 298.9 |
| Cycloheximide | Protein synthesis inhibitor | 0.16 | 0.0140 | >25 | >156.3 |
| Lanatoside C | Cardiotonic agent | 0.19 | 0.0103 | >25 | >131.6 |
| Calcimycin | Antibacterial agent | 0.20 | 0.0165 | 18.10 | 90.5 |
| Digitoxigenin | Cardiotonic agent | 0.29 | 0.0220 | >25 | >86.2 |

<sup>1</sup>DrugBank database (version 5.0) was used for characterizing bioactives <sup>2</sup>50% inhibitory concentration (IC<sub>50</sub>) <sup>3</sup>Standard deviation (SD) of replicated IC<sub>50</sub> values <sup>4</sup>50% cytotoxicity concentration (CC<sub>50</sub>) <sup>5</sup>Therapeutic Index (TI): ratio of CC<sub>50</sub>/IC<sub>50</sub>

Our time-of addition studies demonstrated chloroquine to be effective agains MERS-CoV only if administered no later than 3 hpi (Fig. 2), and this was also the case for ouabain, digitoxin, digoxin, niclosamide, regorafenib, nelfinavir mesylate, ciclesonide, and benidipine hydrochloride (Fig. 2, Supplementary Fig. 2). Important to note, ciclesonide, an immune system suppressor used to treat asthma and allergic rhinitis, was recently shown to inhibit SARS-CoV-2 [29], the cause of COVID-19, and was reported by Japanese medical doctors to have improved pneumonia symptoms in multiple COVID-19 patients [30]. Our data are consistent with previous reports, which indicated that ouabain and other cardiotonic steroids effectively block clathrin-mediated CoV endocytosis [10,27]. In contrast, the minor inhibitory effects we observed for atovaquone, lercanidipine hydrochloride, permethrin, and octocrylene throughout the time-course indicated that these drugs likely act at later stages of the viral life cycle (Supplementary Fig. 2). Notably, our results indicate that lercanidipine hydrochloride and benidipine hydrochloride, both dihydropyridine calcium channel blockers, display different patterns of viral inhibition [31,32]. This observation could be explained by the different channel selectivity of the two drugs: benidipine hydrochloride blocks triple voltage-gated calcium channels, whereas lercanidipine hydrochloride blocks single voltage-gated channels [33-35]. A dendrogram showing the structural relationship of 36 selected inhibitors with anti-MERS-CoV activity is shown in Supplementary Fig. 3.

Combination therapies have become a standard for the treatment for human immunodeficiency virus (HIV) and hepatitis C virus (HCV) infections. They are advantageous over monotherapies due to better antiviral efficacy, reduced toxicity, as well as the ability to prevent the development of viral drug resistance, etc. In this manuscript, we demonstrated that in combination with remdesivir, camostat, and nelfinavir, as well as cepharanthine, have additive and synergistic anti-SARS-CoV-2 effects *in vitro*, respectively. Therefore, our identified drugs, in combination with remdesivir, could be administered to risk group patients prior to the onset of clinical symptoms, thereby reducing the viral load and ideally preventing deterioration of vital signs. Furthermore, depending on the sufficient availability of such drugs, in non-symptomatic COVID-19 patients, such therapies could potentially be employed to reduce the viral load and consequently lower the probability of virus spread in the population, to reduce quarantine periods, etc.

In summary, we identified 12 FDA-approved drugs that could be repurposed for MERS-CoV or potentially COVID-19 therapy, in mono or combination with other drugs. However, further *in vitro* studies are needed to investigate their exact antiviral mechanisms, and to determine potential synergistic effects, in order to prioritize and select drugs for potential use in randomized, double-blind clinical trials, which are mandatory to assess their safe use in humans.

## Supplementary Materials

The following are available in a separate attachment, Supplementary Information, Supplementary Materials and Methods, Figure S1: Example images of MERS-CoV inhibition in Vero cells, Figure S2: Time-of-addition study with additional FDA-approved drugs, Figure S3: Structural relationship between inhibitors, Table S1: Inhibitors identified by HCS with SI >6.

## Author Contributions

All authors contributed to the methodology, validation, formal analysis, investigation, resources, data curation and editing of the manuscript. JYM and MPW conceptualized the study. MPW wrote the manuscript.

## Acknowledgements

We would like to give special thanks to Diana Koo for review and editing of the manuscript.

## Supplementary Information

### Supplementary Material and Methods

#### Reagents

Chloroquine diphosphate (CQ; C6628), chlorpromazine hydrochloride (CPZ; C8138), and cyclosporine A (CsA; C1832) were purchased from Sigma-Aldrich (St. Louis, MO, USA). Lopinavir (LPV; S1380) was purchased from SelleckChem (Houston, TX, USA). Chloroquine was dissolved in Dulbecco’s phosphate-buffered saline (DPBS; Welgene, Gyeongsan, Republic of Korea), and other reagents were dissolved in DMSO for the screening. Anti-MERS-CoV S antibody was purchased from Sino Biological Inc. (Beijing, China). Alexa Fluor 488 goat anti-rabbit IgG (H+L) secondary antibody and Hoechst 33342 were purchased from Molecular Probes/Thermo Fisher Scientific. Paraformaldehyde (PFA) (32% aqueous solution) and normal goat serum were purchased from Electron Microscopy Sciences (Hatfield, PA, USA) and Vector Laboratories, Inc. (Burlingame, CA, USA), respectively.

#### Time-of-addition assays

Twelve FDA-approved drugs were selected after DRC analysis for time-of-addition assays to characterize the mechanism of action. Briefly, Vero cells were seeded at 1.2 × 10^4^ cells per well in Opti-PRO^™^ SFM supplemented with 4 mM L-glutamine and 1× antibiotic-antimycotic solution in black, 96-well, μClear plates (Greiner Bio-One) at 24 h prior to the experiment. Subsequently, cells were infected with MERS-CoV/KOR/2015 at an MOI of 5 at 4°C for 1 h to synchronize the infection, followed by three washes with Dulbecco’s phosphate-buffered saline (DPBS), and transferred to a 37°C incubator. Drugs were added to cells at various time points pre- or post-infection; 1 h prior to or at 0, 1, 2, 3, or 4 hpi. Drugs were added to infected cells at a concentration of 1, 5, or 10 µM depending on the IC_90_ value of the compound. Chloroquine, which was used as an early stage inhibitor control, was used at a concentration of 100 µM. Cells were fixed with 4% PFA at 6 hpi. IFA and image analyses were performed as described above.

### Supplementary Figures

**Fig. S1.**
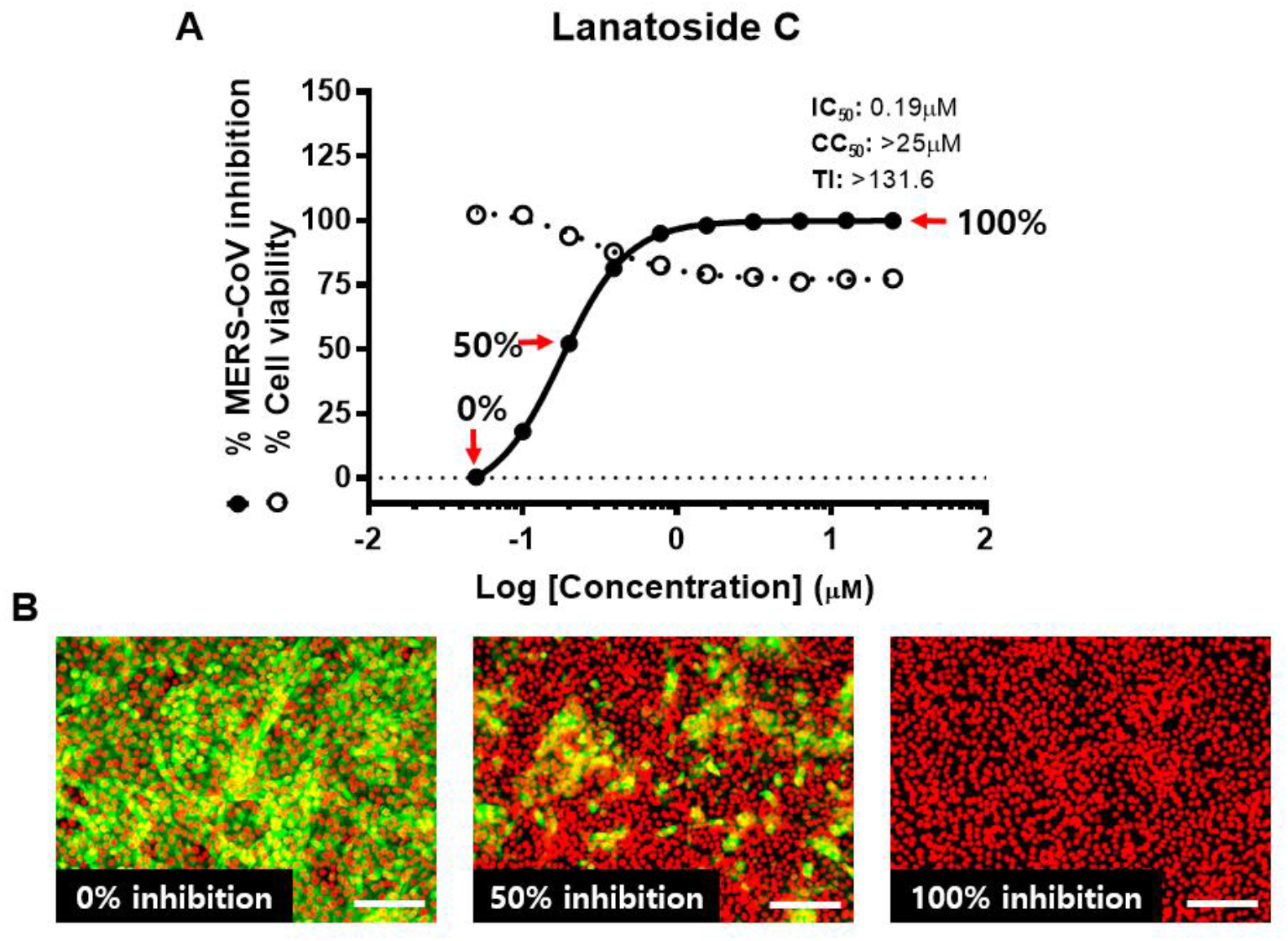
Example images of MERS-CoV inhibition in Vero cells. (A) A representative dose-response curve (DRC) for lanatoside C inhibition of MERS-CoV in Vero cells. (B) HCS was performed using an image-based assay, and compound efficacy was measured by inhibition of MERS-CoV S protein expression. Images depict 0%, 50%, and 100% inhibition, as indicated in the DRC. Scale bar = 100 μm.

**Fig. S2.**
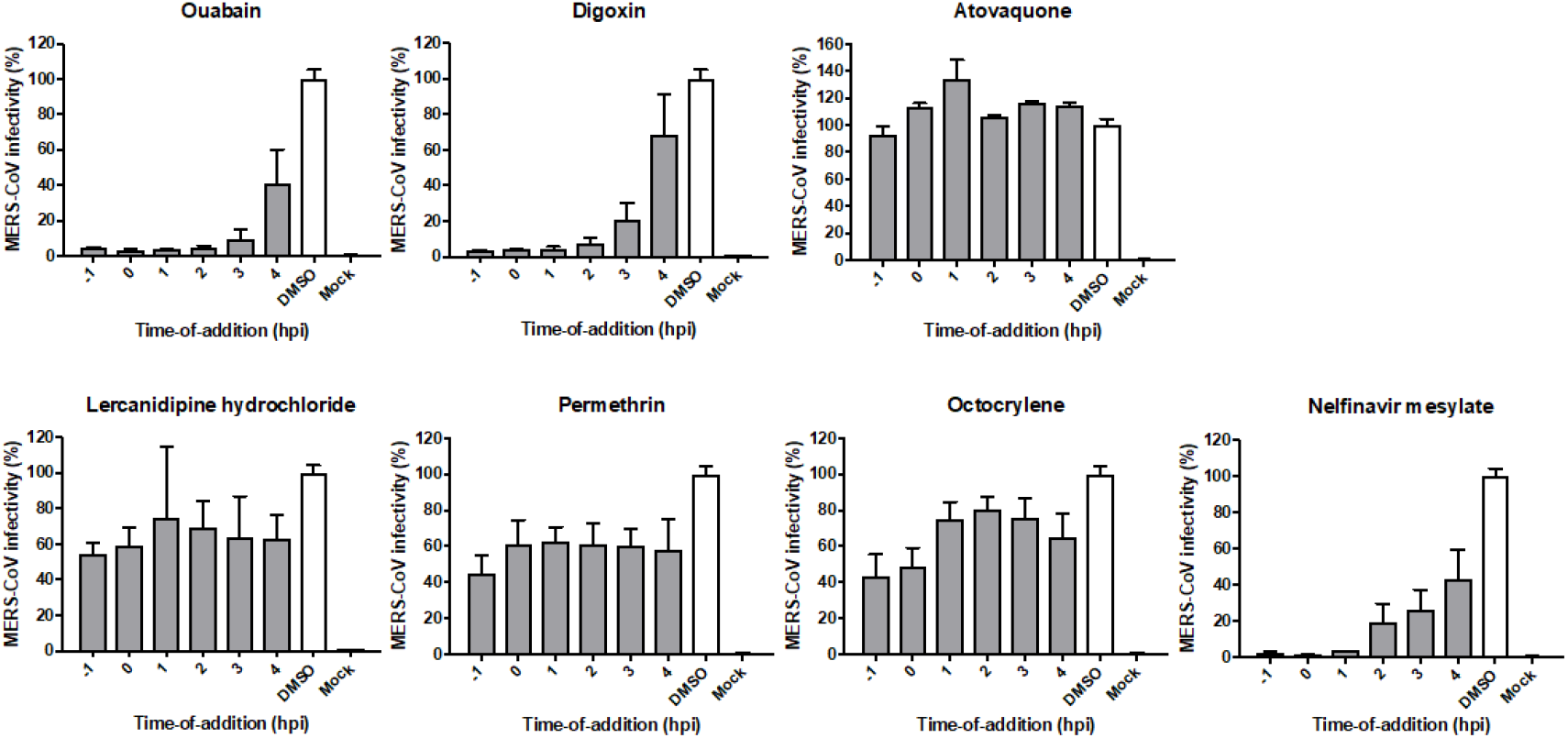
Time-of-addition study with additional FDA-approved drugs. Seven FDA-approved drugs not shown in Fig. 2 were analyzed by time-of-addition experiments. Vero cells were infected with MERS-CoV at a multiplicity of infection of 5, and drugs were administered at six time points pre- or post-infection as indicated. Drugs were used at concentrations above their 90% inhibitory concentration (IC_90_) values.

**Fig. S3.**
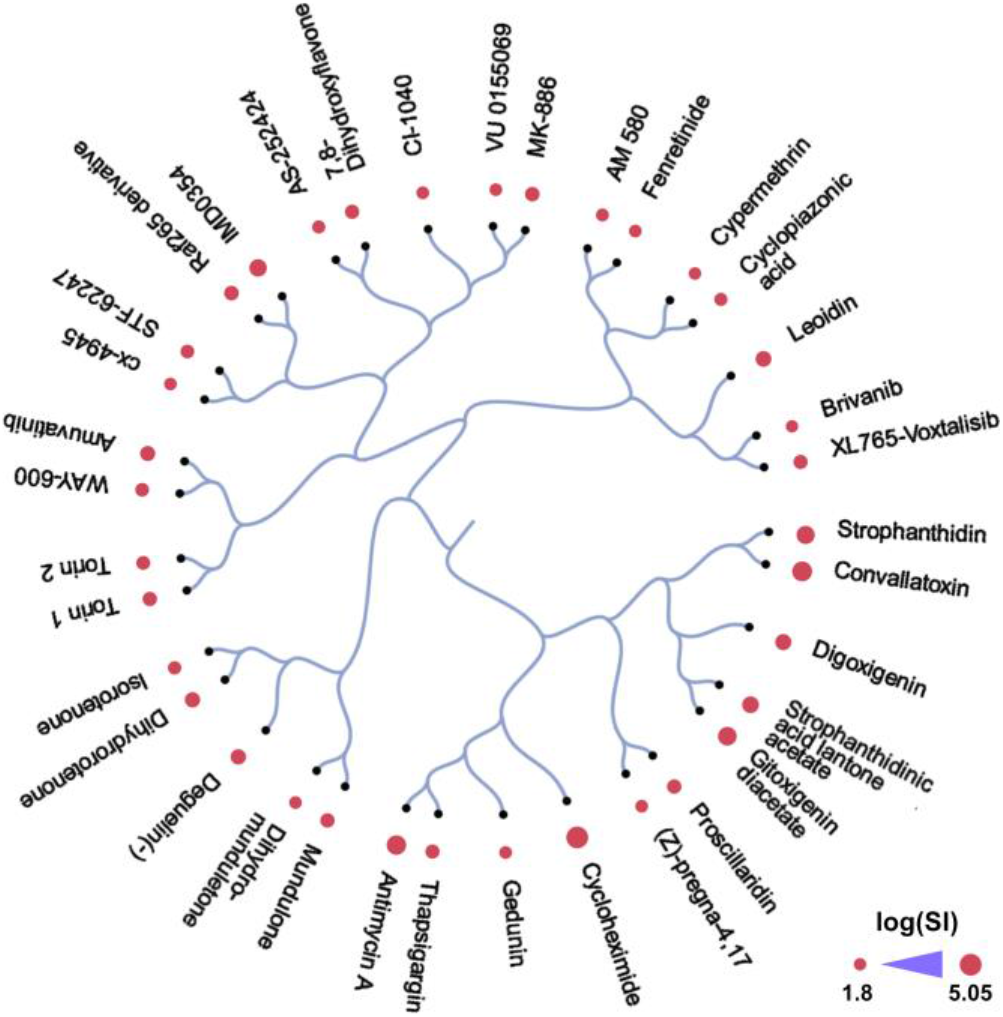
Structural relationship between inhibitors. Dendrogram showing the structural relationship of 36 selected inhibitors with anti-MERS-CoV activity. Compounds were clustered based on their structural similarity calculated by ECPF4 fingerprints and the Tanimoto coefficient [36]. The activity of the compounds was quantified using the log SI values and shown as bubbles. Bubble size corresponds to compounds’ log TI and SI (https://cspade.fimm.fi/).

**Table S1.**
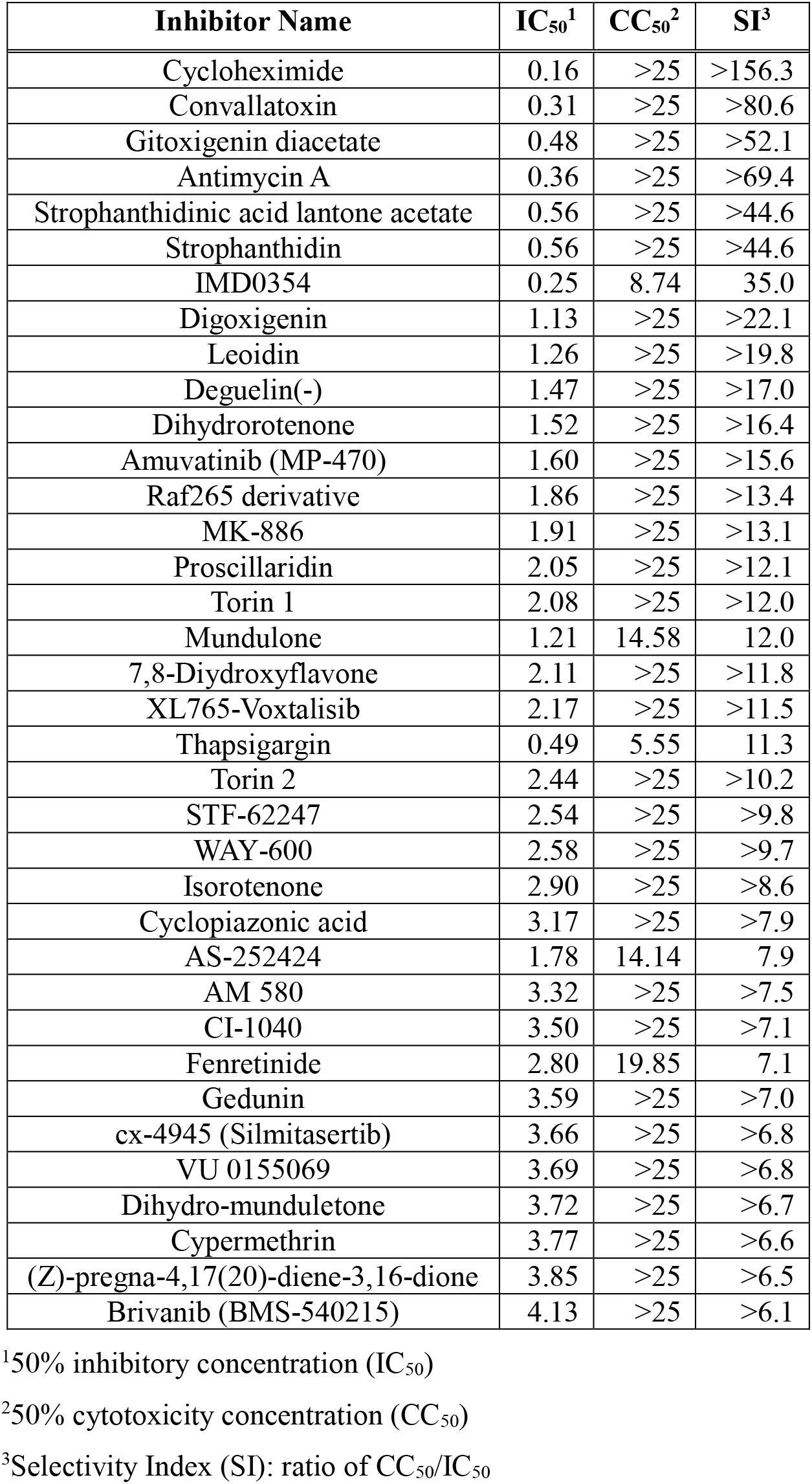
Inhibitors identified by HCS with SI >6.

